# Heterogeneity of *Staphylococcus epidermidis* in prosthetic joint infections: Time to reevaluate microbiological criteria?

**DOI:** 10.1101/2021.03.31.438002

**Authors:** Micael Widerström, Marc Stegger, Anders Johansson, Bharat Kumar Gurram, Anders Rhod Larsen, Lars Wallinder, Helen Edebro, Tor Monsen

## Abstract

Prosthetic joint infection (PJI) is a feared complication after arthroplasty with *Staphylococcus epidermidis* as a major pathogen. One diagnostic criteria for PJI diagnosis is the finding of phenotypically identical organisms based on common laboratory tests in two or more periprosthetic microbial cultures. Because of phenotypical variation within a genetic clone, and clonal variation within a phenotype, the criteria may be ambiguous. Here, we investigate the extent of diversity among coagulase-negative staphylococci in PJI and characterize in detail *S. epidermidis* isolates from these infections.

We performed a retrospective cohort study of 62 consecutive patients with PJI caused by coagulase-negative staphylococci (CoNS) from two hospitals in Northern Sweden. From 16 of the patients, two to nine *S. epidermidis* isolates were available for whole-genome analyses. Hospital-adapted multidrug-resistant genetic clones of *S. epidermidis* were identified in 40/62 (65%) of the PJIs using a combination of analysis by pulsed-field gel electrophoresis and multiple-locus sequence typing. Whole genome sequencing showed presence of multiple sequence types (STs) in seven (7/16, 44%) PJIs. Among isolates of the same ST, within-patient phenotypical variation in antibiotic susceptibility and/or whole-genome antibiotic resistance gene content was frequent (11/16, 69%).

These results highlight the ambiguity of using phenotypical characterization of *S. epidermidis* as diagnostic criteria in PJI. The results call for larger systematic studies to determine the frequency of CoNS diversity in PJIs, the implications of such diversity for microbiological diagnostics, and for the therapeutic outcome in patients.

## INTRODUCTION

Prosthetic joint replacement is one of the most important medical innovations of the 20th century, and it has improved the quality of life for millions of people worldwide by providing pain relief, restoring joint function, mobility and independence (1, 2). In contrast, prosthetic joint infections (PJIs) after joint replacements are devastating complications bringing high hospital costs and increased in–hospital mortality (3). The diagnosis of a PJI and its treatment are both challenging (4). The post-operative infections that normally occur within two years of surgery are not neglectable with an infection rate after hip or knee replacement of between 1-2% (2, 4-9). Register datasets from European countries indicate a significant increase in early revisions for manifest or suspected infection during the last decades (9, 10), likely caused by an aging population and higher levels of obesity in general. These European national quality registers lack data on microbiological etiology, but several investigations have documented that *Staphylococcus aureus* and coagulase–negative staphylococci (CoNS), and in particular *Staphylococcus epidermidis*, accounts for the majority of PJIs (11, 12).

Previous molecular epidemiological studies have revealed clonal spread of hospital-adapted multidrug-resistant *S. epidermidis* (HA–MDRSE) between ward units, hospitals and even countries (13-19). Recently, global spread of HA-MDRSE have been confirmed using whole–genome sequencing (WGS) (20-22), that also highlighted the distinct selection of resistance as indicative of adaption to the hospital environment and the compounds used in the prevention of PJI (23). However, the ramification of the dissemination of HA–MDRSE lineages linked to the increasing incidence of PJI is limited (23-26). This lack of data has historically, in part, been due to the demanding assessment of CoNS in clinical cultures since CoNS constitutes a ubiquitous part of the human skin microbiota, which infers difficulties in distinguishing between isolates from contamination and true infection (27). The current international clinical guidelines for defining a PJI diagnosis reflect this inability since the finding of two positive periprosthetic cultures with phenotypically “identical” or “indistinguishable” organisms is included in the diagnostic set of criteria, with the phenotype being based on common laboratory tests for genus and species identification and antibiograms (28-30). Since phenotypic morphological variation (31), including small colony variants (SCVs) and different antibiogram has been reported in monoclonal CoNS infections (32-34), the term “phenotypically identical organisms” is ambiguous. Here, we investigate the extent of diversity among coagulase-negative staphylococci in PJI and in detail characterize *S. epidermidis* in these infections.

## MATERIALS AND METHODS

### Study population

The study population was recruited from two hospitals in Northern Sweden, Umeå University Hospital (UH), and Östersund County hospital (ÖH), which annually perform approximate 260 and 470 primary hip or knee arthroplasties, respectively, with a reported <2-year infection rate for primary hip arthroplasties of 1.7-2.9% respectively, according to data from the Swedish Arthroplasty Registers (35, 36)

All patients were identified using the laboratory information systems and the presence of CoNS in ≥2 periprosthetic tissue biopsies retrieved from revision surgery in clinically suspected patients with PJI between December 2008 and June 2011. The used clinical workup was to obtain five tissue specimens from different sites of the periprosthetic tissue using new set of sterile instruments for separate specimens, each of which were divided in two: one half was placed in sterile container for direct culturing on blood agar, McLeod agar and anaerobic blood agar and incubated at 37°C for 2 days. The other half was placed in thioglycolate broth and cultured at 37°C for 7 days under anaerobic conditions. Visually negative broths were terminally subcultured on blood agar, McLeod agar and anaerobic agar and incubated for a further 2 days. Diagnosis was based on Infectious Disease Society of America (IDSA, www.idsociety.org) PJI diagnostic criteria ‘identical microorganisms isolated from two or more cultures’ (29) and classified according to when they occurred after implantation: acute, within 1–3 months; delayed, 3 months to 1 year; late, more than 1 years (28). Medical records were reviewed for additional data: Concomitant diseases, previous hospitalization during the last year, intraoperative clinical finding by the surgeon, surgical treatment of the PJI, and outcome at 2-year follow-up.

### Bacterial strains

CoNS cultured from ≥2 perioperative tissue specimens from revision surgery of clinically suspected PJI patients were consecutively collected. Based on difference in morphology, one to two isolates resembling CoNS were picked from each tissue culture for further investigations. The bacterial isolates were stored at –70°C in preservation media (Trypticase Soy Broth, BD Diagnostic Systems, Sparks, MD, USA) until further examination. In total, 131 CoNS isolates from 62 PJIs patients were included for antimicrobial susceptibility testing and pulsed-field gel electrophoresis (PFGE). Only one isolate was available for analysis with PFGE in 40 of the 62 PJIs; 34 *S. epidermidis* PJIs and six non-*S. epidermidis* CoNS PJIs infections (Fig. S1 + Fig. S2). Of the remaining 22 PJIs, 16 included multiple *S. epidermidis* isolates that were available for WGS (Fig. S1). PJIs that had more than one *S. epidermidis* isolate saved were selected for WGS including all available *S. epidermidis* isolates in each patient.

### Identification

CoNS isolates were identified to species level using matrix–assisted laser resorption/ionization time–of–flight mass spectrometry (MALDI-TOF MS), using a Microflex LT (Bruker Daltonik GmbH, Bremen, Germany) and MALDI Biotyper software v3.1 DB7311 (Bruker Daltonik), according to the manufacturer’s instructions. A score >2.0 was required for species identification (37).

### Antibiotic susceptibility testing

Antimicrobial susceptibility testing by disc diffusion was performed according to the recommendations of the European Committee on Antimicrobial Susceptibility Testing (EUCAST, http://www.eucast.org). Briefly, staphylococci were suspended in saline to McFarland 0.5 and inoculated on Mueller-Hinton II Agar (Becton Dickinson, Cockeysville, MD, USA) before application of the antimicrobial discs. Oxacillin resistance was screened using 10 µg cefoxitin disc on Mueller Hinton II Agar supplemented with 2% NaCl.

Constitutive and inducible resistance to clindamycin was determined with the D–shaped disc diffusion method (Oxoid AB, Sweden). Agar plates were incubated at 35°C for 20h before evaluation. The clinical breakpoints were according to EUCAST recommendation (v10.0). Heteroresistance testing for vancomycin was not performed. Multidrug-resistance (MDR) was defined as resistance towards ≥3 antimicrobial classes.

### PFGE and multilocus sequence typing (MLST)

PFGE and MLST were performed as previously described (38). Sequence types (STs) were assigned using the *S. epidermidis* MLST database (https://pubmlst.org/sepidermidis/). PFGE was performed on all isolates (*n*=131). *S. epidermidis* PFGE types that included at least three isolates was further analyzed using MLST.

### Genome sequencing and analyses

Whole-genome sequencing was performed on all 69 *S. epidermidis* isolates available from the 16 PJI patients using Illumina MiSeq and the 300-cycle MiSeq Reagent Kit v3 to generate paired-end 150-bp reads using manufacturer’s instructions with purified DNA using the Qiagen Blood and Tissue kit (Fig S1, Table S1). The generated sequencing data were subjected to quality control using bifrost (https://github.com/ssi-dk/bifrost) to ensure adequate sequencing quality of all isolates. The sequence data was assembled using SPAdes v3.9.0 (39). Raw reads were aligned against the *S. epidermidis* ATCC 12228 reference chromosome (GenBank accession ID CP0222479) for detection of single nucleotide polymorphisms (SNPs) using NASP v1.0.0 (40) after duplicated regions in the reference were removed using NUCmer. NASP was also used to detect intraspecies contamination. All positions with <10-fold coverage or if the variant was present in <90% of the base calls were excluded using GATK (41). The identified SNPs in the core genome was used to infer phylogenetic relationships using PhyML v3 (42) with Smart Model Selection (43).

Resistance mechanisms were detected as previously described (23). Briefly, acquired antimicrobial resistance genes were detected using the curated database used by ResFinder v3.1 (44) to search for gene matches using ABRicate v0.7 (https://github.com/tseemann/abricate) on the assembled genomes using a >80% hit length and >90% sequence identity criteria.

### Data availability

The whole-genome sequence data generated in this study have been submitted to the European Nucleotide Archive under BioProject ID PRJEB44086.

### Statistics

All statistical analyses were performed using the SPSS v24 (SPSS Inc., Chicago, IL, USA) software package. Fisher’s exact test was used to test for association in all two-way tables. A value of p<0.05 was considered significant.

### Research ethics

The study was approved by the Research Ethics Committee (No 2012–477– 31M) of the Faculty of Medicine, Umeå University, Sweden.

## RESULTS

### Clinical characteristics

From December 2008 to June 2011, 62 consecutive patients (34 men and 28 women; median age 68.6 years) with revision or resection arthroplasties due to CoNS-related PJI were identified (Table 1). Thirty–five of 62 (56%) patients were included at UH and 27/62 (44%) at ÖH. The vast majority of patients were diagnosed with hip or knee PJI (59/62, 95%) (Table 1). The distributions were similar between hip or knee revision or centers regarding gender, age, and reasons for primary arthroplasty (data not shown). Early revision or resection surgeries (<3 months after prosthesis implantation) and two–stage exchanges were performed in the majority of patients while debridement, antibiotics and implant retention (DAIR) surgery was used in ∼1/3 (21/62; 34%) of patients. At two-year follow–up, 15/62 (24%) patients had undergone resection, required further surgeries or suppressive antimicrobial treatment and were assessed as failures.

**Table 1.**
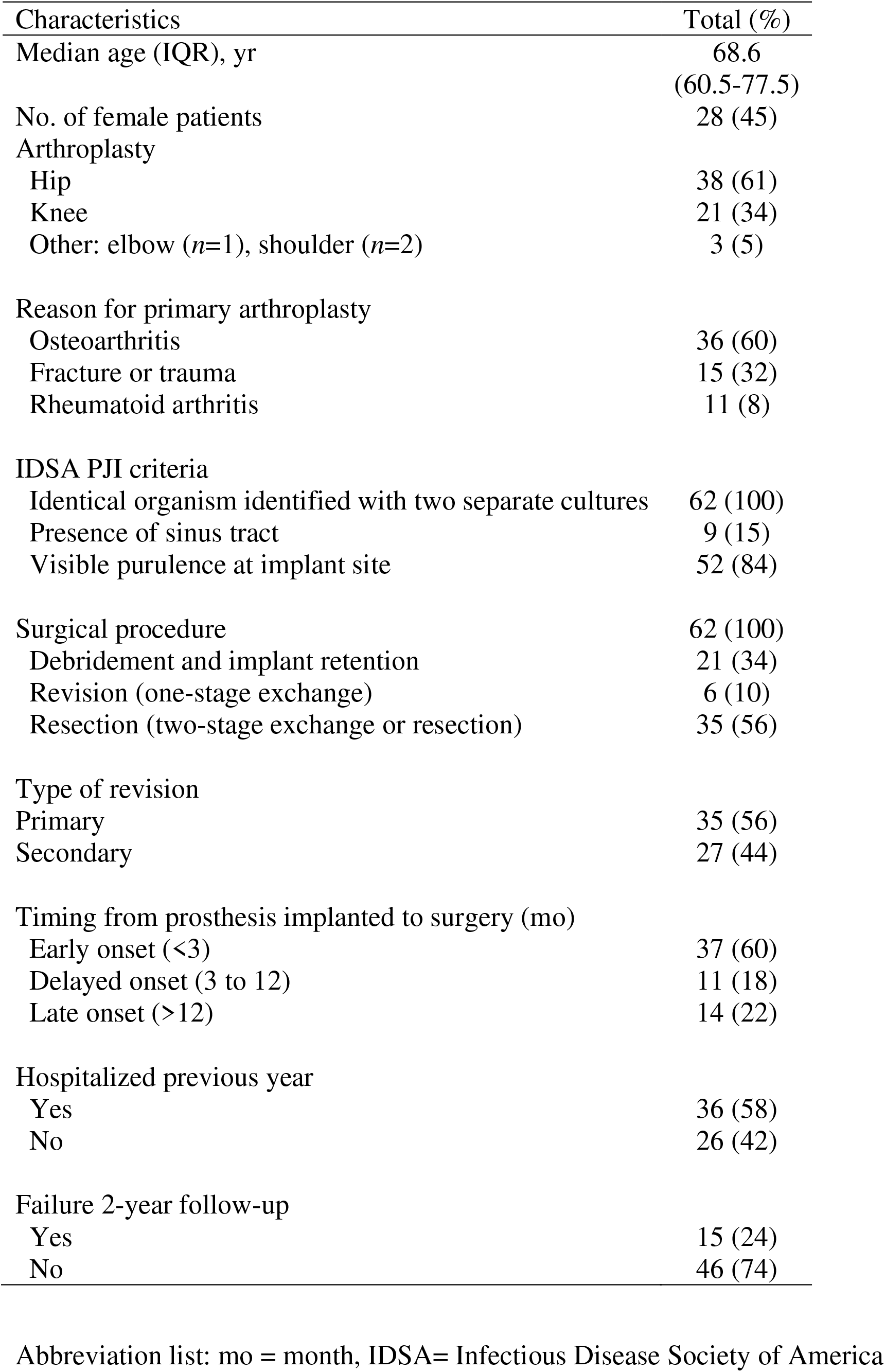
Demographics and clinical characteristics of 62 consecutive patients with prosthetic joint infections due to coagulase-negative staphylococci

### Microbiological findings and genetic analyses by PFGE and MLST

In total, 131 CoNS isolates were available from the 62 patients of PJIs with *S. epidermidis* (*n*=107; 85%), *Staphylococcus capitis* (*n*=11; 8%) and *Staphylococcus hominis* (*n*=8; 4%) as the most frequent species as determined by MALDI-TOF MS (Table 2, Fig. S1). PFGE analysis of all 131 isolates revealed overall genetic clustering by species and subsequent MLST on selected *S. epidermidis* PFGE types revealed two major clusters corresponding to ST215 (*n*=32) and ST2 (*n*=41) (Fig. S2). In addition, two single locus variants (SLVs) of ST215 (ST434) and ST2 (ST188) were identified, the major cluster including SLVs further denoted as the ST215 and ST2 lineages, respectively (Fig. 1). There were 11 patients with genetically indistinguishable isolates using PFGE and MLST (PFGE type A1/ST215), and another seven patients with genetically indistinguishable isolates belonging to PFGE type C/ST2 (Fig. S2). Isolates of the two major lineages ST2 or ST215 were found in 40/52 (77%) of the *S. epidermidis* PJI patients and were equally common among hip and knee PJIs and present throughout the study period at both hospitals (Fig. 1).

**Table 2.**
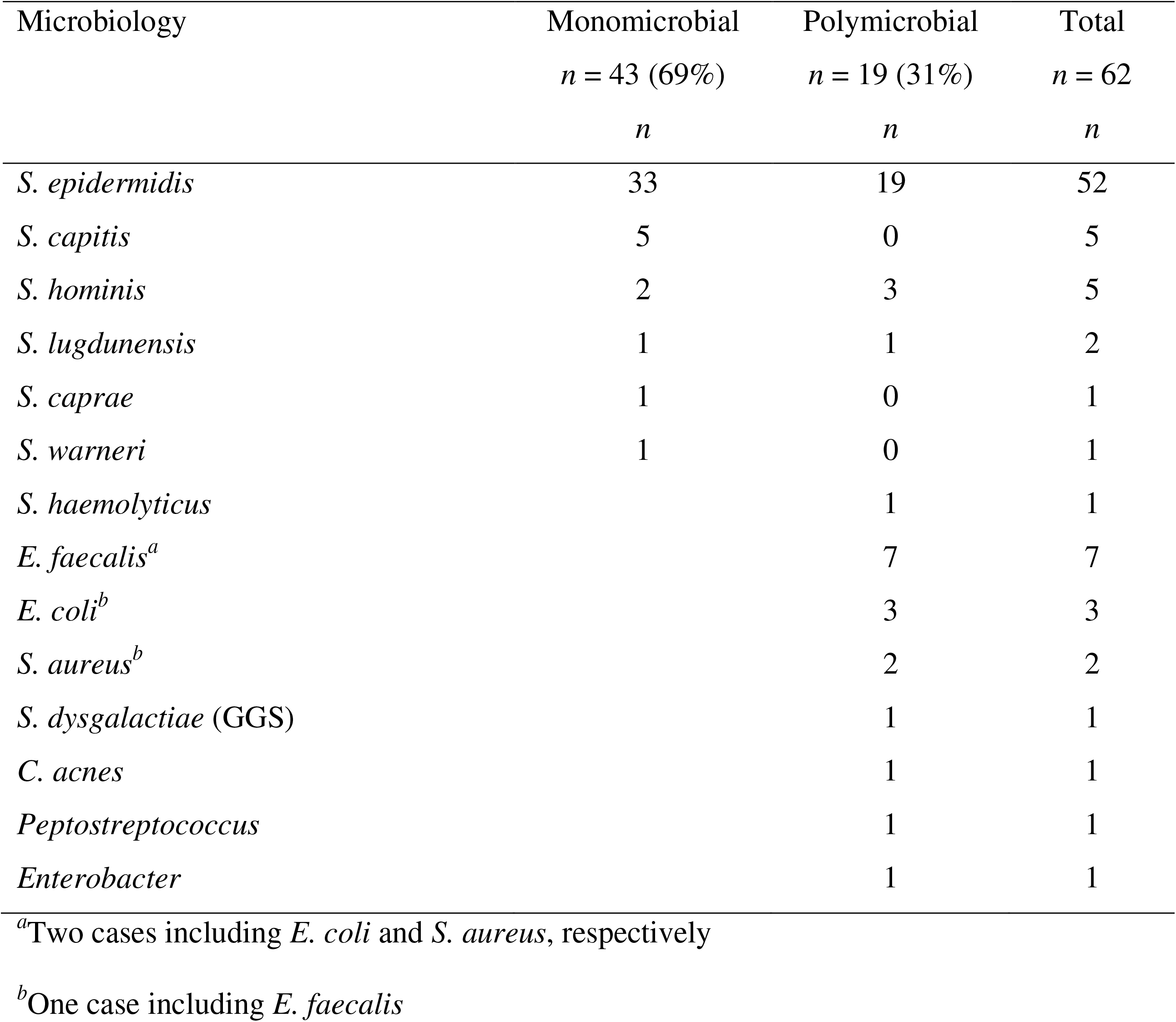
Microbiological characteristics in 62 consecutive patients with prosthetic joint infections due to coagulase-negative staphylococci

**Fig 1:**
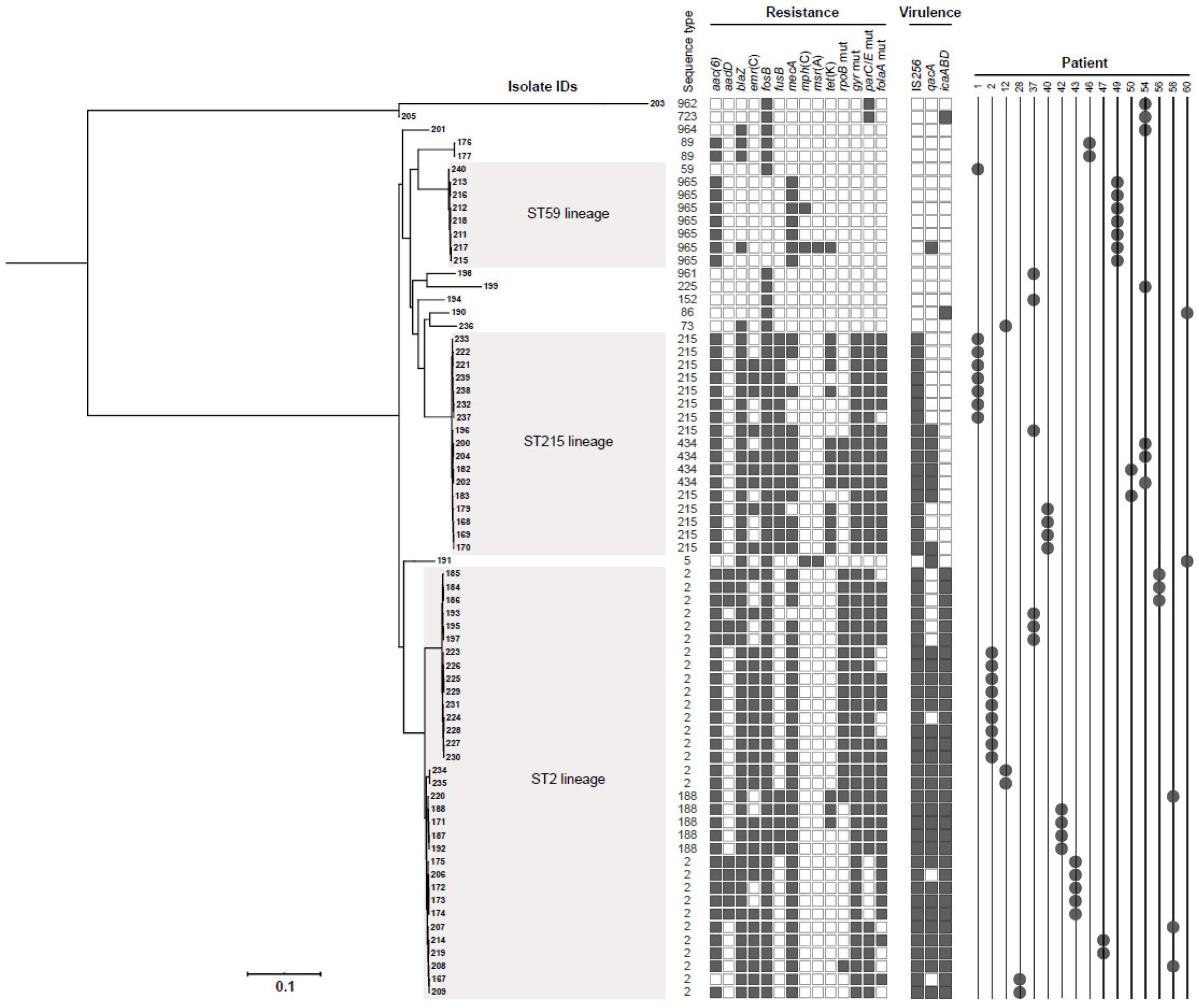
Midpoint-rooted maximum-likelihood phylogeny of 69 *S. epidermidis* PJI isolates from 16 patients. Information on sequence type, genes related to resistance, virulence and biofilm formation is presented as is the patient number. Scale bar indicate substitutions per site. Black blocks represent presence of genes mediating antibiotic resistance or genes previously associated with virulence.

Among *S. epidermidis* isolates, the levels of antimicrobial resistance were as follows: cefoxitin (80%), gentamicin (90%), norfloxacin (79%), trimethoprim-sulfamethoxazole (75%), clindamycin (63%), fusidic acid (42%), and rifampicin 33% (Table S2). No resistance to vancomycin and/or linezolid was detected. Significant differences in antimicrobial susceptibility were identified when comparing the two major genetic clusters (Table S2). All 32 isolates of ST215 lineage exhibited fusidic acid resistance compared to 21% in ST2 lineage (*P*<0.0001). In contrast, rifampicin resistance was significantly more common among isolates in the ST2 lineage than in the ST215 lineage (*P*=0.0002).

### Phenotypic diversity of coagulase-negative staphylococci in PJI

The majority (43/62; 69%) of the PJIs were monomicrobial with *S. epidermidis* identified in 33/43 (77%) of PJI patients followed by *S. capitis* (*n*=5) and *S. hominis* (*n*=2) (Table 2). *S. epidermidis* was identified in all 19 polymicrobial PJIs, most frequently in combination with *Enterococcus faecalis* (*n*=7), *Escherichia coli* (*n*=3) or *S. hominis* (*n*=2), and with similar frequencies in hip and knee revision arthroplasties with 11/37 (30%) and 6/20 (33%), respectively. There was no difference in the distribution of the two major lineages ST215 and ST2 among monomicrobial and polymicrobial PJIs (Table S3). The presence of sinus tract with communication to the joint in PJI patients was reported in only 2/19 (11%) polymicrobial *S. epidermidis* PJIs (*Peptostreptococcus* and *E. faecalis*, respectively). CoNS species, other than *S. epidermidis*, were more common in late PJIs (*P*=0.0004) (data not shown). All ten monomicrobial non-*S. epidermidis* CoNS infections were considered cured at 2-year follow-up compared to 24/33 (73%) monomicrobial *S. epidermidis* PJI and 13/19 (68%) polymicrobial PJIs.

### Whole-genome analyses of *S. epidermidis* in prosthetic joint infections

In 16 of the 62 PJI patients, nine of which were monomicrobial, multiple *S. epidermidis* isolates (*n*=69) in different samples from the same patient were available for WGS (Fig S1, Table 4). Between two to nine isolates per patient were available, and genomic analysis of these identified three major clusters: ST59/ST965 in two PJIs, ST215 lineage in five and ST2 lineage in nine PJIs as illustrated in Fig. 1. Nine of the 16 (56%) patients were infected by a single *S. epidermidis* lineage while seven (44%) patients were infected by between two to five different *S. epidermidis* lineages (Fig. 3). Based on a conserved core genome of 73% (1.83 Mb) across the entire collection of *S. epidermidis* PJI isolates, we found the within-patient genetic diversity among isolates from individual STs ranging from 10 to 107 SNPs, whereas PJI patients infected by multiple STs had 100 to 39,618 SNPs between isolates (Table S1).

**Fig 2.**
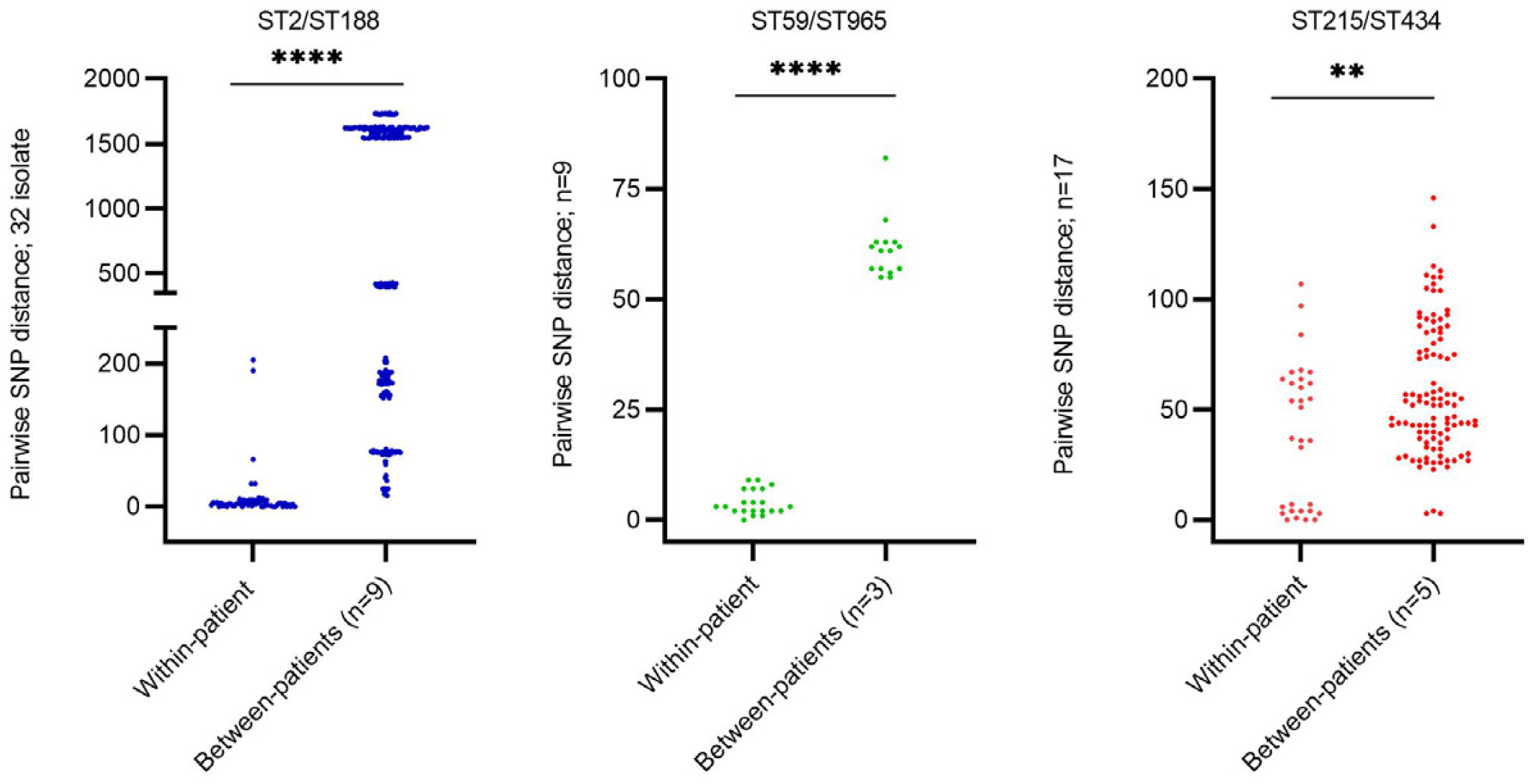
Pairwise SNP within-patient and between-patients SNP distances among *S. epidermidis* isolates belonging to ST2/ST188, ST59/ST965 and ST215/ST434 PJI patients. **** = p<0.0001, ** = 0.002

**Fig 3.**
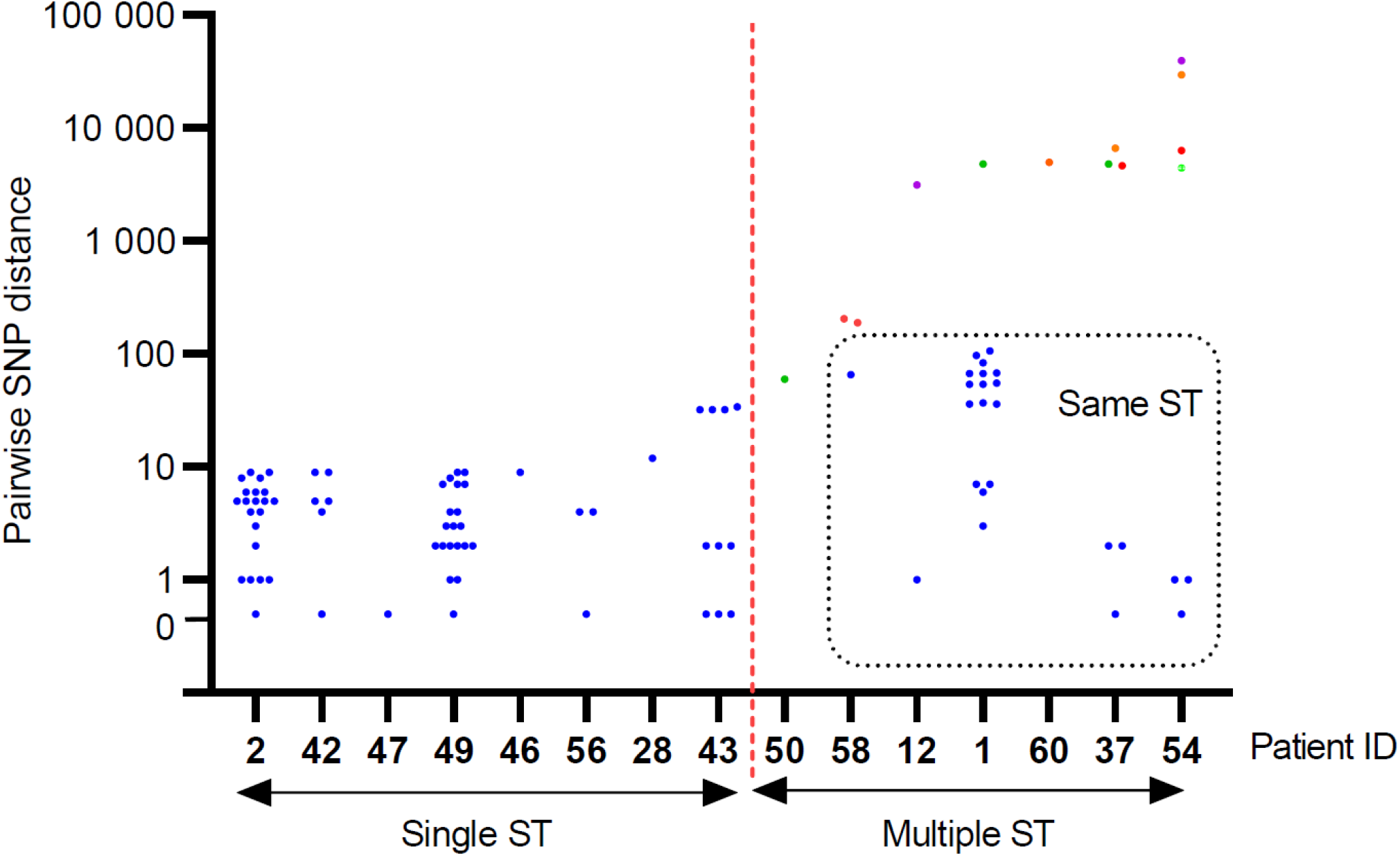
Pairwise within-patient SNP distances among 16 PJI patients where ≥2 *S. epidermidis* isolate were available for analysis. Different colors depict different STs

Analyses of between-patients diversities in the three major STs, revealed mean pairwise SNP distances of 61 for ST59, 58 for ST215, and 1,012 for ST2 (Fig. 2). As should be expected, the within-patient diversity was lower (4, 38 and 13 mean pairwise SNP distances for ST59, ST215 and ST2, respectively) for PJIs with multiple unique isolates of the same lineage. The largest pairwise isolate SNP difference in a patient infected by isolates of a single ST was 107 SNPs (ST215 in patient 1, see Fig. S3). Notably, SNP distances between two isolates of an ST can be similar between-patients and within-patients (Fig. 2). The smallest SNP distance between two ST215 PJI isolates from different patients, that had undergone revision surgery at separate hospitals and two years apart, were 23 SNPs (isolates 238 and 196 from patient 1 and 37, respectively, see Fig 1).

### Virulence and resistance determinants in *S. epidermidis* lineages

There was a high degree of concordance between phenotypic antimicrobial resistance and antimicrobial resistance genes identified by WGS (Table S4). Our analyses showed that some antibiotic resistance genes were also associated with ST types: *fusB* acid *and tet*(K), conferring resistance to fusidic acid and tetracycline respectively, were detected only in ST215/ST434, ST188 and ST59. Likewise, mutations in *gyr*(A), conferring resistance to fluoroquinolones, were detected in isolates belonging to ST2, ST22 and ST215, whereas *rpoB* mutations conferring rifampicin resistance were identified in both ST2 lineages and in sequence type 434, but not in any of the ST215 isolates (Fig. 1, Table S1).

### Within-patient variations in phenotype and genotypic resistance

When we compared multiple *S. epidermidis* isolates collected from individual PJI patients, variation in antibiograms was identified in 13 of the 16 (81%) PJIs (Table 3). The differences in susceptibility ranged from between one to five antimicrobials (Figure 1, Table 4) with colony polymorphism between isolates observed in all patients (data not shown). Variation in antibiotic resistance gene content was also apparent comparing multiple isolates of the same sequence type in one patient (Fig 1, Tabl S1); in patient 1, seven ST215 isolates varied in genetic content regarding *mecA, tet*(K) and *ermC* and *folA* mutations.

**Table 3.** Within-patient polymorphism in phenotypic antimicrobial susceptibility pattern among 16 prosthetic joint infections with ≥2 isolates of *S. epidermidis*

| Patient ID | ST | Isolates<br>(n) | FUS | FOX | GEN | CLI | ERY | NOR | RIF | SXT |
| --- | --- | --- | --- | --- | --- | --- | --- | --- | --- | --- |
| **1** | 59 | 1 | R | S | S | S | S | S | S | S |
|  | 215 | 2 | R | S | R | R | R | R | S | R |
|  | 215 | 1 | R | R | R | S | S | R | S | R |
|  | 215 | 2 | R | S | R | S | S | R | S | R |
|  | 215 | 2 | R | R | R | R | R | R | S | R |
| **2** | 2 | 9 | S | R | R | R | R | R | R | R |
| **12** | 2 | 2 | S | R | R | R | R | R | R | R |
|  | 73 | 1 | S | S | S | S | S | S | S | S |
| **28** | 2 | 1 | S | R | S | R | R | R | S | R |
|  | 2 | 1 | R | R | R | R | R | R | S | R |
| **37** | 2 | 1 | S | S | R | R | R | R | R | R |
|  | 2 | 2 | S | R | R | S | S | R | R | R |
|  | 152 | 1 | S | S | S | S | S | S | S | S |
|  | 215 | 1 | R | R | R | R | R | R | S | R |
|  | 961 | 1 | S | S | S | S | S | S | S | S |
| **40** | 215 | 3 | R | R | R | S | S | R | S | R |
|  | 215 | 1 | R | S | R | R | R | R | S | R |
| **42** | 188 | 3 | R | R | R | R | R | R | S | R |
|  | 188 | 1 | R | R | R | S | S | R | S | R |
| **43** | 2 | 4 | S | R | R | R | R | R | S | R |
|  | 2 | 1 | S | R | R | S | S | R | S | R |
| **46** | 89 | 2 | S | S | R | S | S | S | S | S |
| **47** | 2 | 2 | S | R | R | R | R | R | S | R |
| **49** | 965 | 5 | S | R | R | S | S | S | S | S |
|  | 965 | 1 | R | R | R | S | R | S | S | S |
|  | 965 | 1 | S | R | R | S | R | S | S | S |
| **50** | 215 | 1 | R | R | R | R | R | R | S | R |
|  | 434 | 1 | R | R | R | R | R | R | R | R |
| **54** | 225 | 1 | S | S | S | S | S | S | S | S |
|  | 434 | 3 | R | R | R | R | R | R | R | R |
|  | 723 | 1 | S | S | R | S | S | S | S | S |
|  | 962 | 1 | S | S | S | S | S | S | S | S |
|  | 964 | 1 | S | S | S | S | S | S | S | S |
| **56** | 2 | 2 | S | R | R | S | S | R | R | R |
|  | 2 | 1 | S | R | R | R | R | R | R | R |
| **58** | 2 | 1 | S | R | R | R | R | R | S | R |
|  | 2 | 1 | S | R | R | R | R | R | R | R |
|  | 188 | 1 | R | R | R | R | R | R | R | R |
| **60** | 5 | 1 | S | S | R | S | R | S | S | S |
|  | 86 | 1 | S | S | R | S | S | S | R | S |

**Table 4:** Within-patient phenotypic and genotypic polymorphism among 16 patients with *S. epidermidis* PJI where ≥2 isolate were analyzed using WGS

| Patient ID | Type of implant | Classification<sup>a</sup> | No. of specimens included | No. of *S. epidermidis* isolates | No. of different antibiograms | ST | Polymicrobial | Additional species |
| --- | --- | --- | --- | --- | --- | --- | --- | --- |
| 2 | Hip | Early | 7 | 9 | 1 | 2 | No |  |
| 12 | Hip | Early | 3 | 3 | 2 | 2, 73 | Yes | *S. hominis* |
| 28 | Hip | Early | 2 | 2 | 2 | 2 | No |  |
| 46 | Hip | Early | 2 | 2 | 1 | 89 | No |  |
| 47 | Hip | Delayed | 2 | 2 | 1 | 2 | Yes | *E. faecalis:*<br>*E. coli* |
| 50 | Hip | Delayed | 2 | 2 | 2 | 215, 434 | Yes | *E. faecalis* |
| 56 | Hip | Delayed | 2 | 3 | 2 | 2 | No |  |
| 1 | Hip | Late | 4 | 8 | 5 | 59, 215 | Yes | *S. hominis* |
| 40 | Hip | Late | 4 | 4 | 2 | 215 | Yes | *S. haemolyticus* |
| 43 | Knee | Early | 5 | 5 | 2 | 2 | No |  |
| 54 | Knee | Early | 6 | 7 | 3 | 225, 434, 723,<br>962, 964 | Yes | *S. hominis* |
| 58 | Knee | Early | 3 | 3 | 3 | 2, 188 | No |  |
| 60 | Knee | Early | 2 | 2 | 2 | 5, 86 | Yes | *S. lugdunensis* |
| 42 | Knee | Delayed | 4 | 4 | 2 | 188 | No |  |
| 49 | Knee | Late | 7 | 7 | 3 | 965 | No |  |
| 37 | Shoulder | Late | 5 | 6 | 4 | 2, 152, 215,<br>961 | No |  |

<sup>a</sup>Early = <3 months, Delayed = 3-12 months, Late = >12 months

### Temporal within-patient variations in phenotype and genotypic resistance

*S. epidermidis* isolates from separate patients at different time points were available in four patients (28, 42, 43, 58) (Table S1). In patient 28, two ST2 isolates separated by 518 days, showed acquisition of phenotypic and genotypic gentamicin resistance (*qacA, aac(6’)-aph(2’’))*, phenotypic resistance to fusidic acid (not acquisition of *fusB* or detectable mutation in *fusA*) and reversion of *folA* F99Y mutation. In patient 43, five ST 2 isolates separated by 308 days, had variability of *ermC* content and loss of *qacA*. In patient 58 having two ST 2 and one ST188 isolates, acquisition of *fusB, tet*(K), rifampicin-mutation (S486Y), *folA*-F99Y mutation and loss of *ermC* during treatment with rifampicin and fusidic acid was observed. In patient 42, four ST188 isolates, separated by 162 days, showed variability in content of *emrC* and *tet*(K) and in phenotypic erytromycin/clindamycin resistance.

## DISCUSSION

Here, we investigated the diversity among CoNS in PJI and in detail characterized the *S. epidermidis* isolates in these infections. We found that the within-patient diversity of *S. epidermidis* was considerable with variations in phenotypic and genotypic resistance in the majority (13/16; 81%) of patients, and importantly also when comparing isolates of the same sequence type. In addition, even when considering the inherent difficulty of ruling out the possibility that a single *S. epidermidis* isolate may represent a contamination, *S. epidermidis* isolates belonging to different sequence types were detected in several PJIs (7/16; 44%).

These findings underscore the complexity in assessing whether *S. epidermidis* from multiple cultures in possible PJI cases meet the current criteria for microbiological diagnosis of PJI, i.e. a finding of phenotypically identical organisms in two positive periprosthetic cultures. Hence, with the present guidelines there is a risk that PJIs are incorrectly dismissed as contamination preventing proper microbial diagnosis and treatment.

Before this study, there was limited data on the within-patient genetic diversity of CoNS isolates in PJI (34, 45). Other polyclonal device-related infections have been described, and in addition diversification and evolution of *S. epidermidis* during infection within a patient (45-47). Here, we show that while only a single sequence type was detected in the majority (9/16) of the PJIs, polyclonality was detected in 44% (7/16) of all patients with between two to five different sequence types. Importantly, when at least three *S. epidermidis* isolates were characterized in each PJI, different sequence types were identified in 5/11 (45%) and a difference in antibiograms in almost all patients (10/11, 91%). Obviously, among PJI where only two *S. epidermidis* isolates were available for characterization, polyclonal infection was more rarely detected (2/5 patients, see Table 4). These results are consistent with a recent German study analyzing paired isolates from 55 orthopedic device-related infection cases with a timeframe between 6 to 428 days between isolates, with 6/55 (11%) being assessed as polyclonal (25). The result of the present study suggests that increasing the number of *S. epidermidis* isolates for characterization, and preferably obtained from different tissue specimens is important to determine the diversity among the isolates and reduce the risk of incorrect dismissal as contaminants. Further, the inclusion of more than two isolates will improve the basis for making decisions on antibiotic therapy and for characterizing the clinical condition as a relapse or a reinfection.

We found a clear example of within-patient genetic variation of the ST2 *S. epidermidis* lineage in one of the patients (patient 58). This exemplifies that *S. epidermidis* can adapt to the selective pressures of long-term antimicrobial treatment by acquisition of resistance genes to overcome ongoing rifampicin and fusidic acid treatment. Since polyclonal *S. epidermidis* infections have mainly been reported for device-related infections other than PJI, and considering the reporting of within-patient evolution (45, 47), a more generous definition of “indistinguishable isolates” has been suggested allowing up to two differences in drug susceptibility profiles (48).

Confirming previous data, HA-MDRSE lineages were the cause of the majority of *S. epidermidis* PJI over a period of more than two years in the two hospitals in Northern Sweden (13, 14, 24, 38). The low pair-wise isolate diversity of the ST215 lineage found in two PJIs where isolates were collected more than one year apart in the same hospital (2 SNPs), or separate hospitals (23 SNPs) is supportive for that ST215 is persistent in the hospital setting. This is in contrast to *S. aureus* PJIs where data supports that there is limited hospital-adapted transmission of genetic lineages (49-51). The findings in this study fits with a previously described scenario of a global dissemination of multidrug-resistant lineages of *S. epidermidis* (14, 20, 52, 53). The scenario of hospital-adapted transmission was further corroborated by a recent large PJI study of *S. epidermidis* from Sweden (23). The adaptation of ST2 and ST215 lineages to the hospital environment includes common genomic traits (*IS*256) and and high prevalence of antimicrobial resistance genes even though some lineage dependent differences are evident (20, 23, 54). The primary source of HA-MDRSE lineages and the route of transmission is uncertain. In the subset of 26 PJI patients that had not been hospitalized during the last year, 12 patients were infected by ST215 or ST2 lineages and a majority of these PJI were classified as early infections, suggesting that a history of previous hospitalization is not a prerequisite for acquiring HA-MDRSE PJI. Recent data suggests that current perioperative prevention regimens for PJI selects for MDRSE either from the patients normal flora or by facilitating acquisition from the hospital environment (23).

Polymicrobial infections including *S. epidermidis* were common in the PJIs investigated here and in line with previous data, *E. faecalis* was the most frequent companion microbe (55). In most patients, both *S. epidermidis* and the companion microbe was found in the majority of tissue specimens in each patient, making contamination less likely, even if that possibility cannot be excluded.

The results presented here have practical implications. The finding of within-patient diversity of *S. epidermidis* suggests that the clinical microbiology assessment of a PJI needs to be evaluated (56). Based on our data, characterizing more than two isolates phenotypically and genotypically will improve assessment whether microbiological diagnostic criteria for PJI are met. More than two isolates will also provide more information for deciding on appropriate targeted antibiotic therapy and facilitate the evaluation whether a future infection episode is due to relapse or is a reinfection. The present clinical microbiology methodology for detection of genetic heterogeneity is laborious and expensive but recent advance may change that in the near future. New culture-independent methods that can be applied in clinical laboratories should facilitate rapid assessment of clonality and population structure of *S. epidermidis* communities in PJI (57). Another approach ready to use is culturing of multiple PJI specimens followed by sequencing of multiple microbial isolates as part of PJI routine microbial diagnostics. We think that given the high cost of PJIs, it is realistic to implement routine PJI diagnostics using smaller scale rapid sequencing technology with a turn-around time including bioinformatics of 1-2 days (58)

Limitations of the study include a retrospective cohort design. Although prospective studies are generally preferred we believe that this is not crucial to investigate *S. epidermidis* populations causing PJIs, we note that an earlier study from central Sweden showed that the population structure of *S. epidermidis* remained fairly stable over the last 10 years (23).

Perhaps more important, larger numbers of isolates per patient would have improved the study. Multiple *S. epidermidis* isolates were available for WGS analysis only in a minority of PJIs. With this limitation, the microbiological findings of heterogeneity still underline that the present-day guidelines for PJI diagnosis may be sub-optimal. Lastly and most importantly, it cannot be ruled out that some of the detected and characterized isolates constitute contaminants and are not truly invasive, however, all PJI patients were included consecutively and met IDSA’s criteria for PJI. Further, we used new sets of knife blades for skin incision, subcutaneous incision and new sets of sterile instruments for separate tissue specimens to limit the risk of contamination.

In conclusion, the results outline that the within-patient genetic diversity in *S. epidermidis* isolates was substantial with variation in both antibiotic susceptibility and resistance gene content. The findings highlight the complexity and ambiguity of phenotypical assessment of CoNS isolates from periprosthetic tissue cultures as diagnostic criteria in PJI and calls for larger systematic studies to determine the implications for microbiological diagnosis and the clinical significance of these results for the therapeutic outcome in patients.

## SUPPLEMENTAL MATERIAL

SUPPLEMENTAL FILE 1

## ACKNOWLEDGEMENTS

Preliminary data from this study was presented at ISSSI 2014.

Source of Funding

The project was supported by the following: the Swedish Society of Medicine SLS-24993, SLS-413681; the Research and Development Unit, Jämtland County Council, Sweden; and through a regional agreement between Umeå University and Region Västerbotten. The funders had no role in study design, data collection and interpretation, or the decision to submit the work for publication.

## Figure legends

**Fig S1**. Flowchart depicting the 62 CoNS PJI patients included in the study and the 16 *S. epidermidis* prosthetic joint infections with multiple isolates analyzed by WGS

**Fig. S2**. Dendrogram cluster analysis of the genetic similarity of 131 CoNS isolates using pulsed-field gel electrophoresis (PFGE). The horizontal upper bar indicates genetic similarity (per cent). The dotted lines in the centre of the diagram represent digitalized transformation of the PFGE DNA pattern. The columns to the right present the following: patient ID, PFGE type or non-*S. epidermidis* species and sequence type (ST).

**Fig S3**. Intra-lineage pairwise SNP diversity among *S. epidermidis* STs within each patient with PJI related to time since primary surgery

